# Myricetin protected against Aβ oligomer-induced synaptic impairment, mitochondrial function and oxidative stress in SH-SY5Y cells via ERK1/2/GSK-3β pathways

**DOI:** 10.1101/2023.01.12.523781

**Authors:** Li Wang, Zhi Tang, Yuxin Deng, Yaqian Peng, Yan Xiao, Jianwei Xu, Ruiqing Ni, Xiaolan Qi

## Abstract

Alzheimer’s disease is characterized by abnormal β-amyloid (Aβ) plaque accumulation, tau hyperphosphorylation, reactive oxidative stress, mitochondrial dysfunction and synaptic loss. Myricetin, a dietary flavonoid, has been shown to have neuroprotective effects in vitro and in vivo. Here, we aimed to elucidate the mechanism and pathways involved in myricetin’s protective effect on the toxicity induced by the Aβ42 oligomer. Neuronal SH-SY5Y cells were pretreated with myricetin before incubation with Aβ42 oligomer. The levels of pre- and postsynaptic proteins, mitochondrial division and fusion proteins, glycogen synthase kinase-3 β (GSK-3β) and extracellular regulated kinase (ERK) 1/2 were assessed by Western blotting. Flow cytometry assays for mitochondrial membrane potential (JC1) and reactive oxidative stress, as well immunofluorescence staining for lipid peroxidation (4-HNE) and DNA oxidation (8-OHdG), were performed. We found that myricetin prevented Aβ42 oligomer-induced tau phosphorylation and the reduction in pre/postsynaptic proteins. In addition, myricetin reduced reactive oxygen species generation, lipid peroxidation, and DNA oxidation induced by the Aβ42 oligomer. Moreover, myricetin prevented the Aβ42 oligomer-induced reduction in mitochondrial fusion proteins (mitofusin-1, mitofusin-2), fission protein (dynamin-related protein 1) phosphorylation, and mitochondrial membrane potential via the associated GSK-3β and ERK 1/2 signaling pathways. In conclusion, this study provides new insight into the neuroprotective mechanism of myricetin against Aβ42 oligomer-induced toxicity.

## 1 Introduction

Alzheimer’s disease (AD) is the most common cause of dementia, affecting more than 50 million people worldwide [1, 2]. AD is pathologically characterized by β-amyloid (Aβ) plaques, neurofibrillary tangles formed by hyperphosphorylated tau, and loss of neurons and synapses. In addition, mitochondrial dysfunction and lipid peroxidation are linked to oxidative stress and reactive oxygen species (ROS) in neurotoxicity in the pathological development of AD [3]. Oxidative stress is implicated in neurodegenerative diseases such as AD and diabetes [4]. Acute oxidative stress is closely related to tau phosphorylation [5] and has been shown to stimulate Aβ production and aggregation, which could further lead to tau phosphorylation [6]. In turn, the accumulation of phosphorylated tau in the hippocampus has been previously shown to contribute to oxidative stress and mitochondrial structural and functional changes, forming a vicious cycle [7–10].

Among other signaling pathways, the involvement of aberrant glycogen synthase kinase-3 β (GSK-3β) and extracellular regulated kinase (ERK) signaling pathways in mitochondrial dysfunction and the dysregulation of ROS signaling have been implicated in AD [11, 12]. Aβ monomers form several different intermediate aggregation states, including dimers and trimers, oligomers, protofibrils, fibrils and ultimately Aβ plaques. Aβ oligomers are considered the most neurotoxic form, have been reported in various AD animal models and have been biochemically identified in brain tissue samples from patients with AD [13–15]. Moreover, oligomers and tau have been shown to interact synergistically and amplify the toxic effects on synapses, mitochondria and various downstream pathways [16].

The effects of dietary flavanols and their potential in modulating cognitive function and the risk of AD have gained increasing interest [17–19]. Myricetin (3,3’,4’,5’,5’,7-hexahydroxy-flavone) is a flavonoid found in many natural plants, such as bayberry [20]. Myricetin has been shown to play a critical role in biological pharmacological activities in various fields [21–25]. Myricetin has been shown to reduce Aβ aggregation [26] or Aβ production by restricting the activity of beta-secretase 1 [27, 28] and increasing the level of α-secretase to reduce Aβ production [29]. In addition, myricetin has been shown to restore Aβ-induced mitochondrial impairment [30] and alleviate the response to oxidative stress when redox homeostasis is compromised in the mitochondria [31, 32]. A recent study showed that myricetin prevents high molecular weight Aβ oligomer-induced neurotoxicity through antioxidant effects [33]. Furthermore, myricetin has been shown to eliminate phosphorylated tau and alleviate tau-induced toxicity in neuronal cells [34], potentially by activating autophagy and slowing the liquid liquid phase separation of tau [21]. The effect of myricetin against cognitive impairment has been demonstrated in Aβ42-intoxicated rats [25, 35].

In this study, we hypothesized that myricetin provides a protective mechanism against Aβ42 oligomer (Aβ42O)-induced synaptic toxicity, tau phosphorylation, mitochondrial dysfunction and ROS involving the GSK-3β and ERK1/2 pathways.

## 2 Materials and Methods

### 2.1 Materials, reagents, and antibodies

The list and sources of materials, including chemicals, antibodies, and kits, are described in detail in **Supplemental Table 1** and **Table 2**. Myricetin (purity ≥98.0%) was dissolved first in cell-grade dimethyl sulfoxide (DMSO, 10% v/v) and then further diluted with polyethylene glycol 300 (PEG300, 40%), Tween-80 (5%) and saline (45%). One milligram of Aβ42 freeze-dried powder (purity ≥95.0%) was dissolved in 220 μL hexafluoro-2-propanol, incubated at 37°C for 1 h, transferred to ice for 10 min and air dried in a fume hood at room temperature for 12 h, ensuring that the hexafluoro-2-propanol volatilized completely. The polypeptide film was formed at the bottom and kept at −80□. Nine microliters of cell-grade DMSO was added after the film was fully dissolved. The solution was diluted with 441 μL of F12 medium without phenol red and incubated at 4°C for 24 hours. After centrifugation for 10 min at 14000 r/min, the supernatant was transferred to a new Eppendorf tube and stored at −80°C for future use. Western blotting was first performed to validate the aggregation forms of Aβ using a recombinant anti-Aβ42 antibody (ab180956) (**Supplemental table 1**).

### 2.3 Cell culture, treatment, and extracts

SH-SY5Y cells (well-characterized cellular model of AD) were grown to 80% confluence in 6-well culture plates and maintained in Dulbecco’s modified Eagle medium (DMEM, Gibco, USA) with 10% fetal bovine serum (Gibco, USA) in a humidified incubator with 5% CO_2_ at 37°C. Cells were pretreated with myricetin (5, 10, and 20 μM) for 24 h and then incubated with 10 μM Aβ42O for an additional 48 h.

### 2.3 Cell viability assay

Cell viability was measured using a cell counting kit-8 (CCK-8) (MedChemExpress, USA) as previously described [36, 37]. In brief, a total of 5×10^3^ cells were seeded into 96-well plates. After treatments, 10 μL of CCK-8 reagent and 100 μL of medium were added to each well and incubated for 1 h at 37°C. Next, the absorption (450 nm) of the plate was measured using a microplate reader (Bio-Rad, Hercules, USA).

### 2.4 Western blotting

Next, western blotting was performed to investigate the changes after treatment in the level of presynaptic proteins (SNAP25 and synaptophysin) and postsynaptic proteins (PSD95); phosphorylated-tau (p-tau) and total-tau (t-tau); phosphorylated-ERK (p-ERK) and total-ERK (t-ERK); phosphorylated-GSK3β (p-GSK3β) and total-GSK3β (t-GSK3β) proteins; mitochondrial fission protein (dynamin-related protein 1, Drp1) and fusion proteins (mitofusin-1, Mfn1; mitofusin-2, Mfn2) as previously described [38, 39]. The antibodies used are listed in **Supplementary Table 1**. Total proteins from SH-SY5Y cells were lysed with RIPA lysis buffer (Solarbio, China) containing a protease inhibitor cocktail (1:200, Sigma, USA). Equal amounts of protein per lane (20–50 μg) were loaded onto 4%–12% sodium dodecyl sulfate–polyacrylamide gels (Absin, China) and transferred to polyvinylidene difluoride membranes (Millipore, USA), and the membranes were incubated in 5% skim milk powder (weight/volume) for 2 hours at room temperature. The membranes were incubated with primary antibodies overnight at 4°C and then immersed in goat anti-mouse or goat antirabbit goat anti-mouse IgG (H+L), horseradish peroxidase (HRP), or goat anti-rabbit IgG (H+L), HRP secondary antibodies (Thermo Fisher Scientific, USA). Anti-β-tubulin or anti-β-actin was used as a loading control. Each experiment was repeated at least 3 times. Immunoreactive bands were visualized using a ChemiDoc MP imaging system (Bio-Rad, USA). Band density was analysed by ImageJ software (NIH, USA).

### 2.6 Mitochondrial membrane potential detection by flow cytometry

The mitochondrial membrane potential assay (JC-1 detection kit, Beyotime, China) was implemented to quantify the level of polarization/depolarization of the mitochondrial membrane after treatment with myricetin and Aβ42O in SH-SY5Y cells. SH-SY5Y cells were treated with 10 μM carbonyl cyanide m-chlorophenyl hydrazone for 30 min as a positive control. Cells were digested and collected in 1.5 mL Eppendorf tubes after three gentle washes with phosphate-buffered saline (PBS). A total of 1×10^6^ cells were added to 0.5 mL 1×JC-1 dye (diluted by JC-1 staining buffer and purified water). and incubated at 37°C for 30 min. After washing the JC-1 staining dye with 1×JC-1 buffer three times, 10^4^ cells from each group were examined by using a FACSVerse^™^ flow cytometer (BD Biosciences, USA) at excitation wavelengths of 488 and 633 nm. Fluorescence data were recorded and analysed by using Flow Jo software (BD Bioscience, USA). Each experiment was repeated at least 3 times independently.

### 2.5 ROS detection by flow cytometry

To detect the level of intracellular ROS in SH-SY5Y cells after treatment with myricetin and Aβ42O, a flow cytometry assay with the fluorescent probe 2,7-dichlorofluorescein diacetate (DCFH-DA, Beyotime, China) was performed. At a density of 2×10^5^, SH-SY5Y cells were inoculated in 6-well cell culture plates as described earlier [40]. Cells were maintained in DMEM containing 10 μM DCFH-DA at 37°C for 30 min. The staining solution containing DCFH-DA was then removed. The cells were then gently washed three times with DMEM. The cell suspension was loaded into a flow-specific tube and detected by flow cytometry at excitation and emission wavelengths of 485 and 538 nm in sequence. Fluorescence was measured by BD FACSVerse^™^ flow cytometry (BD Biosciences, USA). Flow Jo software (BD Bioscience, USA) was used to analyse the data. Each experiment was repeated at least 3 times independently.

### 2.7 Measurement of markers of lipid peroxidation and DNA damage

The levels of 8-hydroxy-2’-deoxyguanosine (8-OHdG) and 4-hydroxynonenal (4-HNE) produced by lipid peroxidation as DNA damage markers after treatment with myricetin and Aβ42O were evaluated. SH-SY5Y cells were grown on polylysine-coated coverslips and fixed with 4% paraformaldehyde in PBS pH 7.4 for 30 min at room temperature. Cells were permeabilized by treatment with 0.1% Triton X-100 (Solarbio, China) for 5 min and then incubated with primary antibodies against 8-OHdG (1:200) and 4-HNE (1:200) overnight at 4°C. After washing with PBS pH 7.4, fluorescent secondary antibody conjugated with Alexa Fluor 488 (1:200, ThermoFisher Scientific, USA) or Alexa Fluor 546 (1:200, ThermoFisher Scientific, USA) was added and incubated for 1 h. The sections were then mounted with vector medium containing 4’,6-diamidino-2-phenylindole (DAPI) Fluoromount-G (Vector Laboratories, USA). All images were examined on an Olympus confocal microscope (Olympus, Japan) with a 40× objective and a constant exposure time for each marker in all sections analysed. The experiments were repeated three times. A total of 100 cells from each group were analysed. Each experiment was repeated at least 3 times independently.

### 2.8 Statistical analysis

All analyses were performed with SPSS software, version 25.0 (IBM, USA) and GraphPad Prism version 9.0 (GraphPad, USA). Data are presented as mean ± standard error (SD) (n=3 or 6). One hundred cells from each group were analysed. Statistical significance was determined by one-way ANOVA, followed by Tukey’s multiple comparison test. A p value <0.05 was considered statistically significant.

## 3 RESULTS

### 3.1 Dose response analysis in myricetin and Aβ42O

First, we determined the cytotoxicity and optimal dosage of myricetin (1-200μM) and Aβ42O (1-60μM) using the CCK-8 cell viability test in SH-SY5Y cells. No loss of cell viability was observed in SH-SY5Y cells treated for 24h with 1μM to 15μM myricetin compared to controls treated without myricetin. A dose response in the SH-SY5Y cell viability loss was observed with treatment of 20-200μM myricetin (20-70%) (treated vs. control group) (**Figs. 1a, b**). Thus, a concentration range of 5, 10, and 20μM myricetin was selected in the following investigations. Oligomeric Aβ42 (estimated mass 10-72kDa) was the major species, including 16-mers, tetramers, tremers, dimers and monomers (**Fig. 1c**). Aβ42O reduced the cell viability from 7.5 to 60 μM by 10-30% (**Fig. 1d**). Thus, a concentration range of 10 μM Aβ42O was selected in the following investigations.

**Figure 1.**
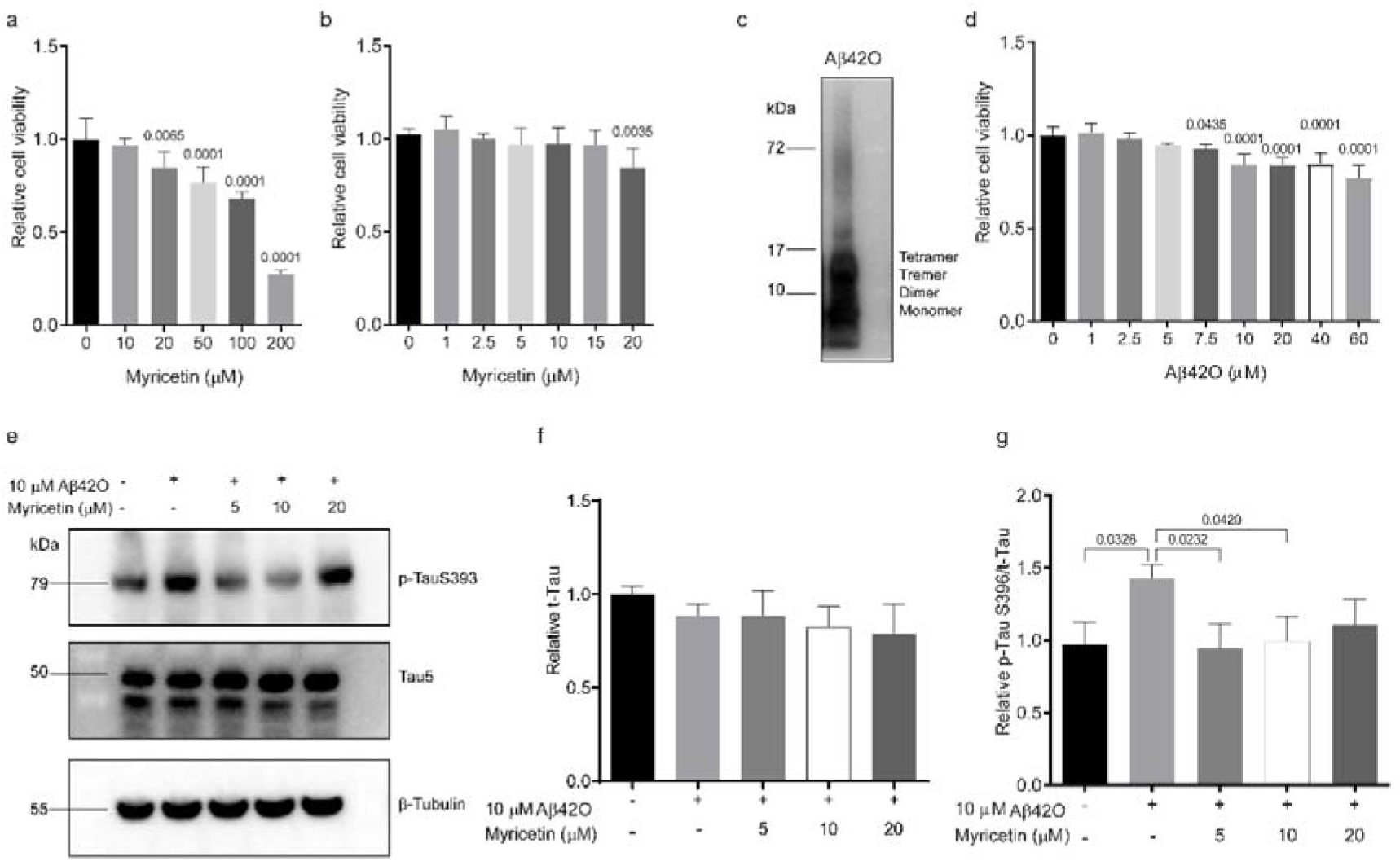
Myricetin decreased the Aβ42O-induced tau phosphorylation in SH-SY5Y cells. **(a, b, d)** Dose-response for cell viability test for myricetin and Aβ42O was assessed by using the CCK-8 assay. SH-SY5Y cells were treated without and with myricetin and Aβ42O at different concentration gradients for 24 h (n=6). Data are presented as the mean ± SD. One-way ANOVA, Dunnett’s multiple comparison. (c) Representative blots of Aβ42O. **(e**) Representative blots of TauS396 and total tau (Tau5); (**f**) Tau5 levels remained stable with different treatments. Values were normalized to the control. **(g**) Myricetin decreased the Aβ42O-induced increase in the p-TauS396/Tau5 ratio. Values were normalized to the control. Data are presented as the mean ± SD, one-way ANOVA, Tukey’s multiple comparison.

### 3.2 Myricetin prevented Aβ42O-induced tau phosphorylation in SH-SY5Y cells

Next, we assessed the influence of myricetin (5-20 μM) pretreatment on the Aβ42O (10 μM)-induced effect on tau phosphorylation in SH-SY5Y cells. We found no change in the t-tau level in all groups (**Figs. 1e, f**). Treatment with Aβ42O increased the relative expression of p-tau/t-tau (Taus396/Tau5) by 43% (p=0.0328, control vs. Aβ42O group, **Figs. 1e, g**), which was prevented by myricetin (5 μM p=0.0232, 10 μM p=0.0420, myricetin+Aβ42O vs. Aβ42O group) (**Figs. 1e, g**).

### 3.2 Myricetin ameliorated Aβ42O-induced synaptic impairment in SH-SY5Y cells

Next, we assessed the influence of pretreatment with myricetin (5-20 μM) on the Aβ42O (10 μM)-induced effect on synaptic impairment in SH-SY5Y cells. Treatment with Aβ42O reduced the relative expression of presynaptic proteins SNAP25 (by 32%) and synaptophysin (by 23%) and postsynaptic proteins PSD95 (by 17%, p=0.0146 control vs. Aβ42O group, **Figs. 2a-c**). Pretreatment with myricetin completely prevented this Aβ42O-induced reduction in presynaptic SNAP25 expression (10 μM, p=0.0496, myricetin+Aβ42O vs. Aβ42O group) (**Figs. 2a, b**) and synaptophysin expression in SH-SY5Y cells (5 μM p=0.0071, myricetin+Aβ42O vs. Aβ42O group) (**Figs. 2a, c**). Pretreatment with myricetin (5 and 10 μM) ameliorated the Aβ42O (10 μM)-induced reduction in PSD95 expression in SH-SY5Y cells (**Figs. 2a, d**).

**Figure 2.**
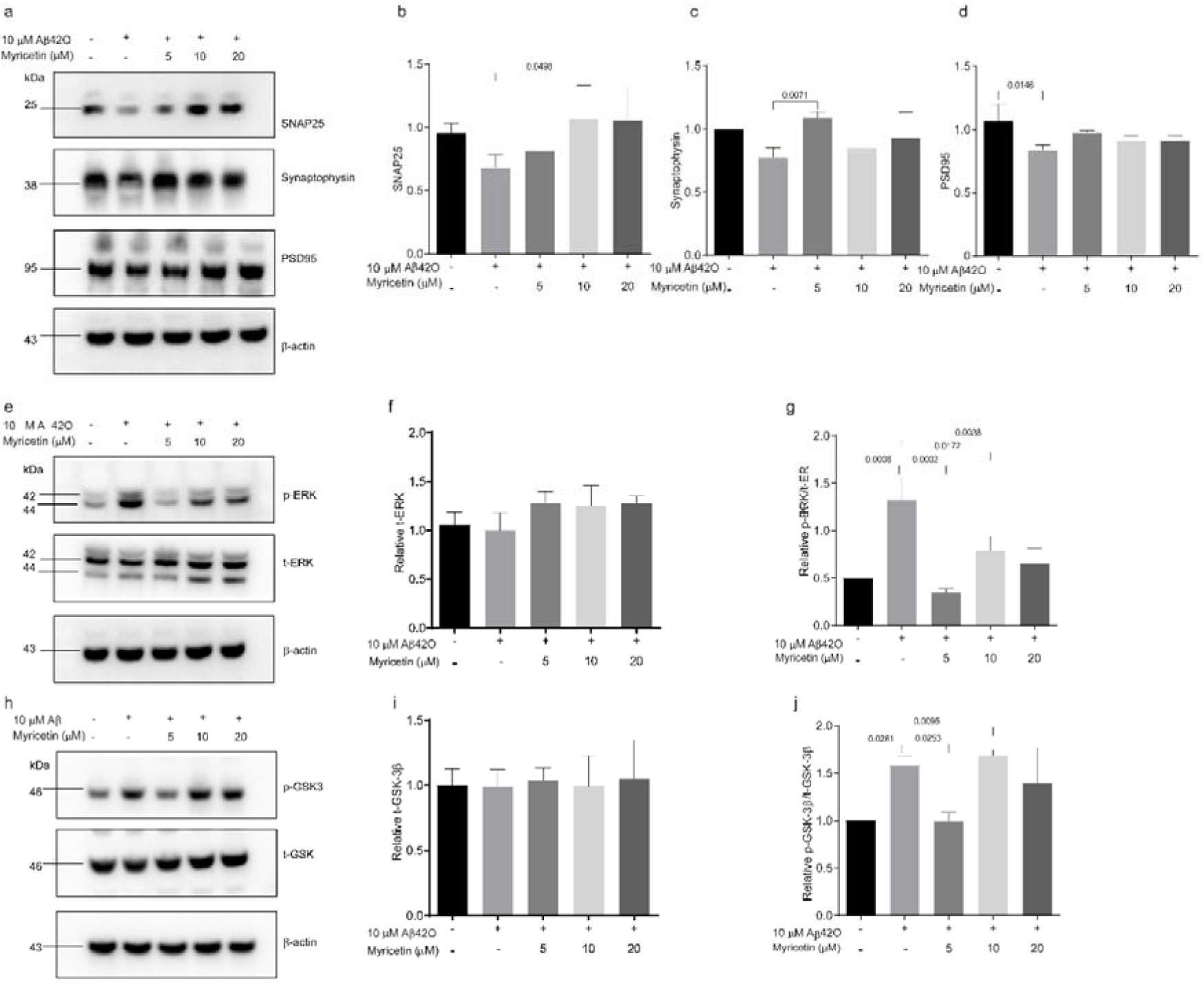
Myricetin enhanced the Aβ42O-induced reduction in synaptic proteins and attenuated p-GSK-3β and p-ERK1/2 expression in SH-SY5Y cells. (**a-d**) Representative blots and quantification of the expression of presynaptic SNAP25, synaptophysin, and postsynaptic PSD95. (**e-g**) Representative blots and quantification of t-GSK-3β and p-GSK-3β/t-GSK-3β. (**h-j**) Representative blots and quantification of t-ERK1/2 and p-ERK1/2 (n=3). Values were normalized to the control. Data are presented as the mean ± SD, one-way ANOVA, Tukey’s multiple comparison.

### 3.3 Myricetin prevented Aβ42O-induced phosphorylation in the ERK1/2 and GSK-3β pathways in SH-SY5Y cells

We hypothesized that the ERK1/2 and GSK-3β signaling pathways were involved in the neuroprotective effects of myricetin on Aβ42O-induced alterations. We assessed the influence of myricetin pretreatment on Aβ42O’s effect on p-ERK1/2, t-ERK1/2, p-GSK-3β and t-GSK-3β in SH-SY5Y cells. We found no change in the t-ERK1/2 or t-GSK-3β level in all groups (**Figs. 2e, f, h, i**). Treatment with Aβ42O (10 μM) elevated the relative expression of p-ERK1/2/t-ERK1/2 and p-GSK-3β/t-GSK-3β (by 165% p=0.0008, 58%, p=0.0261 control vs. Aβ42O group). Pretreatment with myricetin prevented this Aβ42O (10 μM)-induced increase in p-ERK1/2 (5 μM p=0.0002, 10 μM p=0.0172, 20 μM p=0.0038, control vs. Aβ42O group) and p-GSK-3β at 5 μM (p=0.0253, control vs. Aβ42O group) but not at 10 μM or 20 μM (**Figs. 2e, g, h, j**).

### 3.4 Myricetin restored Aβ42O-induced mitochondrial fission protein, mitochondrial membrane potential reduction and DRP1 phosphorylation in SH-SY5Y cells

Next, we assessed the influence of myricetin pretreatment on the effect of Aβ42O on mitochondrial function in SH-SY5Y cells. Treatment with Aβ42O (10 μM) reduced the relative expression of the mitochondrial fission proteins Mfn1 (by 43%) and Mfn2 (by 30%) (p=0.0309 and p=0.0004, control vs. Aβ42O group). Myricetin pretreatment restored the Aβ42O-induced reduction in Mfn1 expression (10 μM p=0.0353, 20 μM p=0.0198, myricetin+ Aβ42O vs. Aβ42O group) and Mfn2 expression (5 μM p=0.008, 10 μM p=0.003, 20 μM p=0.0027, myricetin+Aβ42O vs. Aβ42O group) (**Figs. 3a-c**). We found no change in the t-DRP1 level in all groups but increased the relative expression of p-DRP1/t-DRP1 by 33% (p=0.0190, control vs. Aβ42O group). Pretreatment with myricetin (5 μM but not 10 μM) prevented this Aβ42O-induced increase in p-DRP1/t-DRP1 expression in SH-SY5Y cells (p=0.0386, myricetin+ Aβ42O vs. Aβ42O group) (**Figs. 3a, d, e**).

**Figure 3.**
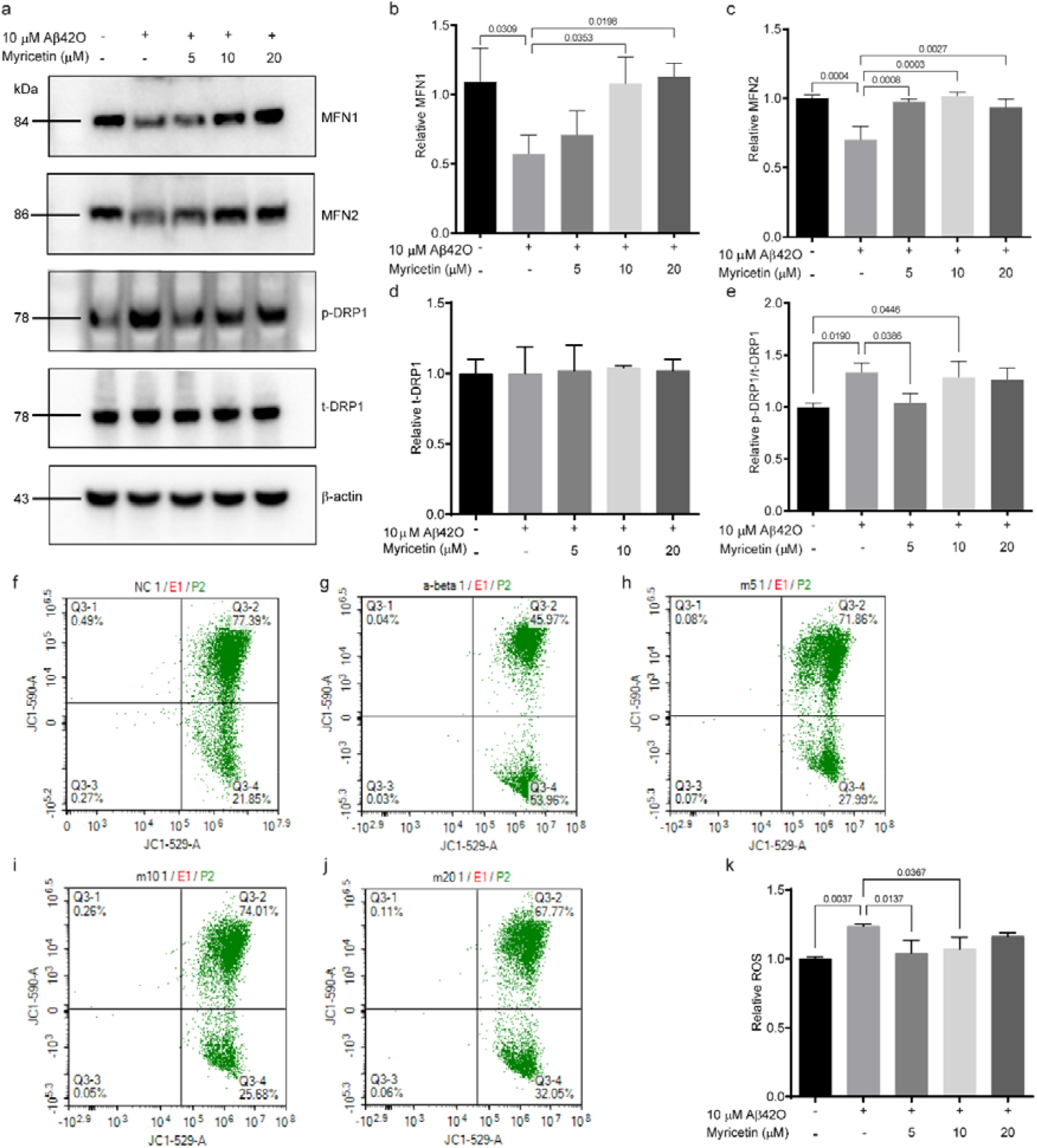
Myricetin regulated Aβ42O-induced mitochondrial fusion protein reduction, mitochondrial division protein phosphorylation, and ΔΨm depolarization in SH-SY5Y cells. (**a-e**) Representative blots and quantification of mitochondrial fusion protein (Mfn1, Mfn2) and mitochondrial division (t-Drp1 and p-Drp1/t-Drp1) (n=3). Values were normalized to the control. (**f-k**) Analysis of ΔΨm with JC-1 staining by flow cytometry (n=3). (f) Control, (g) 10 μM Aβ42O treatment, (h) 10 μM Aβ42O + 5μM myricetin treatment, (i) 10 μM Aβ42O + 10 μM myricetin treatment, (j) 10μM Aβ42O + 20 μM myricetin treatment, (k) Quantification of green/red fluorescence intensity (normalized to control). One-way ANOVA, Tukey’s multiple comparison.

We examined the effect of myricetin on Aβ42O-induced alterations in membrane potential ΔΨm by using flow cytometry. When Ψm is diminished, JC-1 aggregates (emitting red fluorescence) are converted to the monomer state, emitting green fluorescence. We found that treatment using Aβ42O (10 μM) increased the ratio of green/red fluorescence, indicative of the monomer/J-aggregate ratio, by 272% (p<0.0001, control vs. Aβ42O group, **Figs. 3f, k**). Pretreatment with myricetin (5-20 μM) prevented this Aβ42O-induced increase in the monomer/J-aggregate ratio in SH-SY5Y cells (p<0.0001 for all three conditions, myricetin+ Aβ42O vs. Aβ42O group) (**Figs. 3f-k**).

### 3.6 Myricetin blocked Aβ42O-induced ROS production, lipid peroxidation, and DNA oxidation in SH-SY5Y cells

Next, we evaluated the effects of myricetin on Aβ42O-induced ROS generation in SH-SY5Y cells by using flow cytometry with a DCFH-DA assay. We found that treatment with Aβ42O (10 μM) increased ROS by 24% (p=0.0037, control vs. Aβ42O group), which was completely blocked by myricetin (5 μM p=0.0137, 10 μM 0,0367, myricetin+ Aβ42O vs. Aβ42O group) (**Figs. 4a-f**). We found that treatment with Aβ42O (10 μM) increased lipid peroxidation, as indicated by 8-OHdG fluorescence intensity, by 61% (p<0.0001, control vs. Aβ42O group). Pretreatment with myricetin (5. 10, 20 μM) prevented this Aβ42O (10 μM)-induced increase in 8-OHdG fluorescence intensity in SH-SY5Y cells (p<0.0001 for all groups, myricetin+ Aβ42O vs. Aβ42O group) (**Figs. 4g, h**). Furthermore, we found that treatment with Aβ42O (10 μM) increased lipid peroxidation, as indicated by 4-HNE fluorescence intensity by 116% (p<0.0001, control vs. Aβ42O group). Pretreatment with myricetin (5. 10, 20 μM) prevented this Aβ42O (10 μM)-induced increase in 4-HNE fluorescence intensity in SH-SY5Y cells (p<0.0001 for all groups, myricetin+ Aβ42O vs. Aβ42O group) (**Figs. 4i, j**).

**Figure 4.**
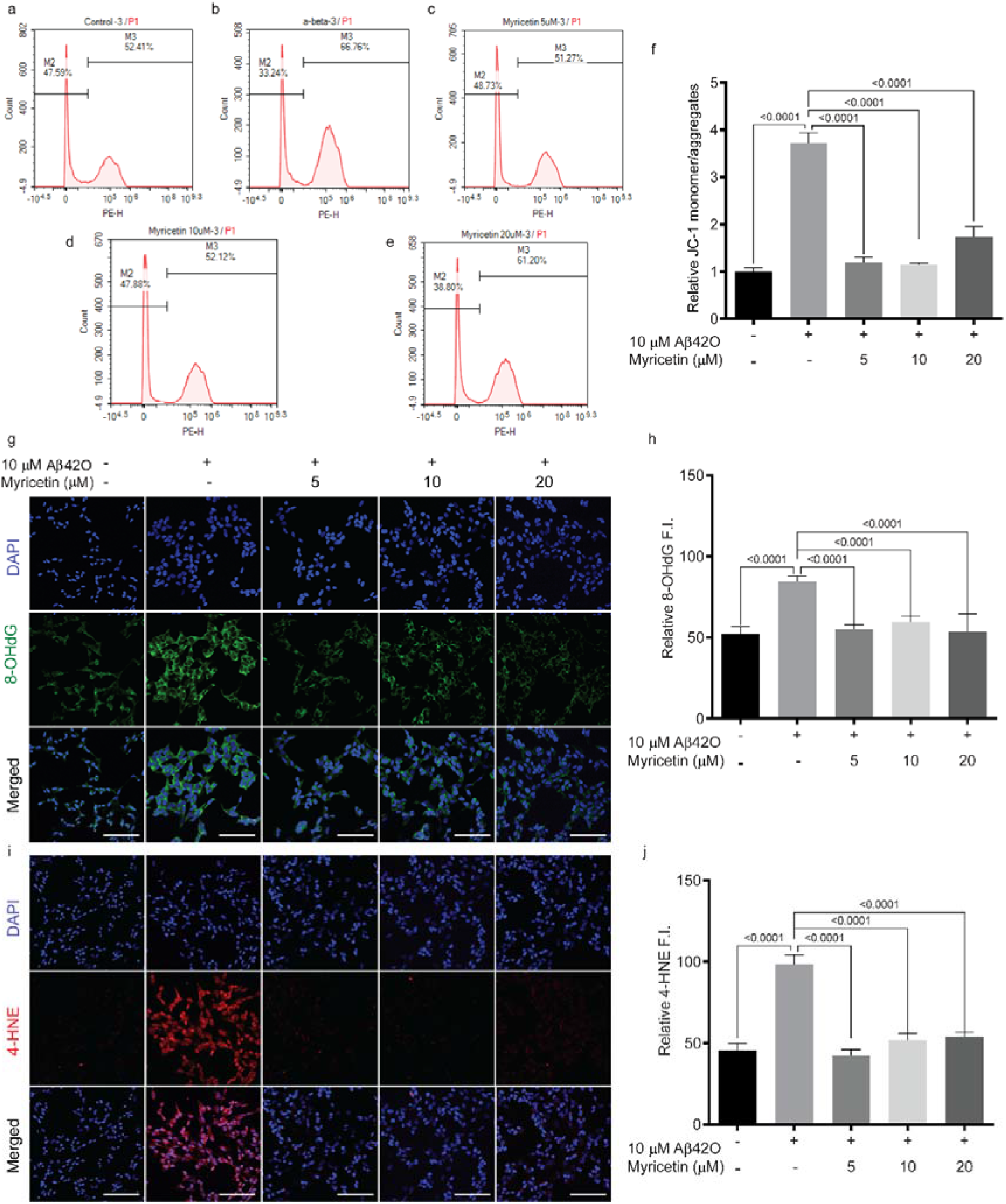
Myricetin reversed the Aβ42O-induced increase in ROS production, lipid peroxidation and DNA oxidation in SH-SY5Y cells. (**a-f**) Analysis of ROS with DCFH-DA by flow cytometry, (a) Control, (b) 10 μM Aβ42O treatment, (c) 10 μM Aβ42O + 5 μM myricetin treatment, (d) 10 μM Aβ42O+10 μM myricetin treatment, (e) 10 μM Aβ42O + 20μM myricetin treatment, (f) Quantification of ROS (normalized to control). (**g, h**) Representative confocal images and quantification of the immunoreactivity of 8-OHdG. Nuclei are counterstained with DAPI; (**i, j**) Representative confocal images and quantification of the immunoreactivity of 4-HNE. Nuclei were counterstained with DAPI; scale bar=100 μm (g, i). A total of 100 cells from each group were analysed. One-way ANOVA, Tukey’s multiple comparison.

## 4 Discussion

Here, we demonstrated the protective effects of myricetin on Aβ42O-induced toxicity in tau hyperphosphorylation, synaptic and mitochondrial impairment, ROS production and oxidation. Myricetin’s effect against Aβ42O involved the inhibition of hyperphosphorylation of GSK-3β and the ERK1/2 signaling pathway.

We found that myricetin restores Aβ42O-induced synaptic impairment by increasing the reduced expression of presynaptic proteins (SNAP25 and synaptophysin) and the postsynaptic protein PSD95 and tau phosphorylation, which has not been demonstrated previously. Our results are consistent with previous reports that Aβ oligomers have been shown to decrease synaptic protein expression indicative of synaptic function [13–15, 41] and promote the level of tau phosphorylation [16]. Myricetin has been previously shown to prevent Aβ oligomerization and reduce neuronal plasticity (long-term potentiation and longterm depression) by site-specific binding to Aβ [42].

We observed that myricetin restored the Aβ42O-induced reduction in mitochondrial membrane potential, expression of mitochondrial fusion proteins (Mfn1, Mfn2), and phosphorylation of mitochondrial fission protein (p-Drp1). Several studies have shown that Aβ reduces the expression of Mfn1, Mfn2 and Drp1 in animal models of AD [8, 43, 44]. Our results are consistent with recent studies showing that myricetin prevented high molecular weight Aβ42 oligomer-induced neurotoxicity through antioxidant effects in the cell membranes and mitochondria of HT-22 cells [33] and by restoring Aβ-induced mitochondrial impairments in N2a-SW cells [30]. However, our result is different from the finding of one previous report of the stable expression levels of various fission/fusion-mediating proteins, such as Mfn1 and Mfn2, and the phosphorylation of Drp1 after 1–6 h of Aβ treatment [47].

The potential reason is the duration of treatment of Aβ and forms of Aβ oligomers in different studies. A recent study indicated the involvement of mitochondrial fission and mitophagy in Aβ oligomer-induced synaptotoxicity [45]. Phosphorylation of Drp1 has been reported to activate Drp1-mediated fission, and translocation of Drp1 is important in the initiation of fission [46].

We found that myricetin prevented the Aβ42O-induced increased generation of intracellular ROS, lipid peroxidation (4-HNE), and DNA oxidation (8-OHdG). This finding is consistent with previous reports that myricetin reduced lipid peroxidation and DNA oxidation in human periodontal ligament stem cells [48]. ROS overproduction is known to interfere with the equilibrium between oxidants and antioxidant capacity, leading to extensive damage to subcellular structures, mitochondrial dysfunction, lipid peroxidation, and DNA damage [3, 49]. The reciprocal association between mitochondrial fusion proteins and ROS in Aβ-treated neurons has been reported previously [47, 50].

Moreover, we found that phosphorylation of ERK1/2 and GSK-3β was involved in the protective effect of myricetin against Aβ42O in SH-SY5Y cells. This observation is in line with previous findings on myricetin’s effect on the ERK1/2 and GSK-3β pathways [39, 51]. ERK1/2 plays a central role in oxidative stress in AD, has been shown to be involved in tau phosphorylation [52, 53] and is a potential therapeutic target in various neurodegenerative disorders [54]. GSK-3β, a downstream protein kinase of AKT, plays an important role in tau hyperphosphorylation, regulation of hippocampal neurogenesis and synaptic plasticity and is involved in mitochondrial function [55–57]. GSK-3β also functions in DNA repair by phosphorylating DNA repair factors and influencing their binding to chromatin [56].

There are several limitations in our study. 1) We observed that myricetin showed a reduction in cell viability at a higher concentration (20 μM), which is quite close to the effective dosage of 5 μM. 2) Here, we only examined the involvement of GSK-3β and the ERK1/2 pathway. However, different pathways, such as the BMP-2/Smad and ERK/JNK/p38 mitogen-activated protein kinase signaling pathways and the mTOR pathway ATG5-dependent autophagy pathway, have also been implicated for myricetin [21, 48]. 3) Further in vivo studies in animal models of AD with cerebral Aβ accumulation are needed.

## 5. Conclusions

We show that myricetin inhibits Aβ42O-induced tau phosphorylation, impairment in pre- and postsynaptic proteins, mitochondrial function, ROS generation, lipid peroxidation, and DNA oxidation. Furthermore, hyperphosphorylation of the GSK-3β and ERK1/2 signaling pathways was involved in the protective effect of myricetin against Aβ42O. These findings provide new insights into the protective mechanism of myricetin against Aβ42O-induced toxicity.

## Authors’ Contributions

ZT, XLQ, and RN contributed to the conception and design of the study. LW, YXD, and YQP performed the experiments. LW, YX, and YXD contributed to data collection and analysis. LW and ZT interpreted the data. XLQ, LW, XLQ, RN and ZT wrote the manuscript. All authors approved the manuscript before submission.

## References

1. 2022 Alzheimer’s disease facts and figures. Alzheimer’s & dementia: the journal of the Alzheimer’s Association 2022, 18(4):700–789.

2. Knopman DS, Amieva H, Petersen RC, Chételat G, Holtzman DM, Hyman BT, Nixon RA, Jones DT: Alzheimer disease. Nature reviews Disease primers 2021, 7(1):33.

3. Park MW, Cha HW, Kim J, Kim JH, Yang H, Yoon S, Boonpraman N, Yi SS, Yoo ID, Moon JS: NOX4 promotes ferroptosis of astrocytes by oxidative stress-induced lipid peroxidation via the impairment of mitochondrial metabolism in Alzheimer’s diseases. Redox biology 2021, 41:101947.

4. Zhang J, Liu L, Zhang Y, Yuan Y, Miao Z, Lu K, Zhang X, Ni R, Zhang H, Zhao Y et al: ChemR23 signaling ameliorates cognitive impairments in diabetic mice via dampening oxidative stress and NLRP3 inflammasome activation. Redox Biology 2022, 58:102554.

5. Alavi Naini SM, Soussi-Yanicostas N: Tau Hyperphosphorylation and Oxidative Stress, a Critical Vicious Circle in Neurodegenerative Tauopathies? Oxid Med Cell Longev 2015, 2015:151979.

6. Petrozziello T, Bordt EA, Mills AN, Kim SE, Sapp E, Devlin BA, Obeng-Marnu AA, Farhan SMK, Amaral AC, Dujardin S et al: Targeting Tau Mitigates Mitochondrial Fragmentation and Oxidative Stress in Amyotrophic Lateral Sclerosis. Molecular neurobiology 2022, 59(1):683–702.

7. Denechaud M, Geurs S, Comptdaer T, Bégard S, Garcia-Núñez A, Pechereau LA, Bouillet T, Vermeiren Y, De Deyn PP, Perbet R et al: Tau promotes oxidative stress-associated cycling neurons in S phase as a pro-survival mechanism: Possible implication for Alzheimer’s disease. Prog Neurobiol 2022:102386.

8. Kandimalla R, Manczak M, Yin X, Wang R, Reddy PH: Hippocampal phosphorylated tau induced cognitive decline, dendritic spine loss and mitochondrial abnormalities in a mouse model of Alzheimer’s disease. Human molecular genetics 2018, 27(1):30–40.

9. Xu M, Huang H, Mo X, Zhu Y, Chen X, Li X, Peng X, Xu Z, Chen L, Rong S et al: Quercetin-3-O-Glucuronide Alleviates Cognitive Deficit and Toxicity in Aβ(1-42) - Induced AD-Like Mice and SH-SY5Y Cells. Molecular nutrition & food research 2021, 65(6):e2000660.

10. Zhang GH, Pare RB, Chin KL, Qian YH: Tβ4 ameliorates oxidative damage and apoptosis through ERK/MAPK and 5-HT1A signaling pathway in Aβ insulted SH-SY5Y cells. Life sciences 2021:120178.

11. Lauretti E, Dincer O, Praticò D: Glycogen synthase kinase-3 signaling in Alzheimer’s disease. Biochimica et biophysica acta Molecular cell research 2020, 1867(5):118664.

12. Khezri MR, Yousefi K, Esmaeili A, Ghasemnejad-Berenji M: The Role of ERK1/2 Pathway in the Pathophysiology of Alzheimer’s Disease: An Overview and Update on New Developments. Cellular and molecular neurobiology 2022.

13. Benilova I, Karran E, De Strooper B: The toxic Aβ oligomer and Alzheimer’s disease: an emperor in need of clothes. Nature Neuroscience 2012, 15(3):349–357.

14. Shankar GM, Li S, Mehta TH, Garcia-Munoz A, Shepardson NE, Smith I, Brett FM, Farrell MA, Rowan MJ, Lemere CA et al: Amyloid-β protein dimers isolated directly from Alzheimer’s brains impair synaptic plasticity and memory. Nature Medicine 2008, 14(8):837–842.

15. Brinkmalm G, Hong W, Wang Z, Liu W, O’Malley TT, Sun X, Frosch MP, Selkoe DJ, Portelius E, Zetterberg H et al: Identification of neurotoxic cross-linked amyloid-β dimers in the Alzheimer’s brain. Brain 2019, 142(5):1441–1457.

16. Busche MA, Hyman BT: Synergy between amyloid-β and tau in Alzheimer’s disease. Nat Neurosci 2020, 23(10):1183–1193.

17. Holland TM, Agarwal P, Wang Y, Leurgans SE, Bennett DA, Booth SL, Morris MC: Dietary flavonols and risk of Alzheimer dementia. Neurology 2020, 94(16):e1749–e1756.

18. Holland TM, Agarwal P, Wang Y, Dhana K, Leurgans SE, Shea K, Booth SL, Rajan K, Schneider JA, Barnes LL: Association of Dietary Intake of Flavonols With Changes in Global Cognition and Several Cognitive Abilities. Neurology 2022.

19. Wang Q, Dong X, Zhang R, Zhao C: Flavonoids with Potential Anti-Amyloidogenic Effects as Therapeutic Drugs for Treating Alzheimer’s Disease. J Alzheimers Dis 2021, 84(2):505–533.

20. Song X, Tan L, Wang M, Ren C, Guo C, Yang B, Ren Y, Cao Z, Li Y, Pei J: Myricetin: A review of the most recent research. Biomedicine & pharmacotherapy = Biomedecine & pharmacotherapie 2021, 134:111017.

21. Dai B, Zhong T, Chen ZX, Chen W, Zhang N, Liu XL, Wang LQ, Chen J, Liang Y: Myricetin slows liquid-liquid phase separation of Tau and activates ATG5-dependent autophagy to suppress Tau toxicity. The Journal of biological chemistry 2021, 297(4):101222.

22. Pan X, Chen T, Zhang Z, Chen X, Chen C, Chen L, Wang X, Ying X: Activation of Nrf2/HO-1 signal with Myricetin for attenuating ECM degradation in human chondrocytes and ameliorating the murine osteoarthritis. International immunopharmacology 2019, 75:105742.

23. Yang W, Yang M, Tian Y, Jiang Q, Loor JJ, Cao J, Wang S, Gao C, Fan W, Zhang B et al: Effect of Myricetin on Lipid Metabolism in Primary Calf Hepatocytes Challenged with Long-Chain Fatty Acids. Metabolites 2022, 12(11).

24. Sun WL, Li XY, Dou HY, Wang XD, Li JD, Shen L, Ji HF: Myricetin supplementation decreases hepatic lipid synthesis and inflammation by modulating gut microbiota. Cell reports 2021, 36(9):109641.

25. Ramezani M, Darbandi N, Khodagholi F, Hashemi A: Myricetin protects hippocampal CA3 pyramidal neurons and improves learning and memory impairments in rats with Alzheimer’s disease. Neural regeneration research 2016, 11(12):1976–1980.

26. Hamaguchi T, Ono K, Murase A, Yamada M: Phenolic compounds prevent Alzheimer’s pathology through different effects on the amyloid-beta aggregation pathway. Am J Pathol 2009, 175(6):2557–2565.

27. Shimmyo Y, Kihara T, Akaike A, Niidome T, Sugimoto H: Multifunction of myricetin on A beta: neuroprotection via a conformational change of A beta and reduction of A beta via the interference of secretases. Journal of neuroscience research 2008, 86(2):368–377.

28. Chakraborty S, Kumar S, Basu S: Conformational transition in the substrate binding domain of β-secretase exploited by NMA and its implication in inhibitor recognition: BACE1-myricetin a case study. Neurochemistry international 2011, 58(8):914–923.

29. Feng J, Wang JX, Du YH, Liu Y, Zhang W, Chen JF, Liu YJ, Zheng M, Wang KJ, He GQ: Dihydromyricetin inhibits microglial activation and neuroinflammation by suppressing NLRP3 inflammasome activation in APP/PS1 transgenic mice. CNS neuroscience & therapeutics 2018, 24(12):1207–1218.

30. Yao X, Zhang J, Lu Y, Deng Y, Zhao R, Xiao S: Myricetin Restores Aβ-Induced Mitochondrial Impairments in N2a-SW Cells. ACS Chem Neurosci 2022, 13(4):454–463.

31. Mendes V, Vilaça R, de Freitas V, Ferreira PM, Mateus N, Costa V: Effect of myricetin, pyrogallol, and phloroglucinol on yeast resistance to oxidative stress. Oxidative medicine and cellular longevity 2015, 2015:782504.

32. Rehman MU, Rather IA: Myricetin Abrogates Cisplatin-Induced Oxidative Stress, Inflammatory Response, and Goblet Cell Disintegration in Colon of Wistar Rats. Plants (Basel, Switzerland) 2019, 9(1).

33. Kimura AM, Tsuji M, Yasumoto T, Mori Y, Oguchi T, Tsuji Y, Umino M, Umino A, Nishikawa T, Nakamura S et al: Myricetin prevents high molecular weight Aβ(1-42) oligomer-induced neurotoxicity through antioxidant effects in cell membranes and mitochondria. Free Radic Biol Med 2021, 171:232–244.

34. Pluta R, Januszewski S, Czuczwar SJ: Myricetin as a Promising Molecule for the Treatment of Post-Ischemic Brain Neurodegeneration. Nutrients 2021, 13(2).

35. Halder T, Patel B, Acharya N: Design and optimization of myricetin encapsulated nanostructured lipid carriers: In-vivo assessment against cognitive impairment in amyloid beta ((1-42)) intoxicated rats. Life Sci 2022, 297:120479.

36. Chen Q, Lai C, Chen F, Ding Y, Zhou Y, Su S, Ni R, Tang Z: Emodin Protects SH-SY5Y Cells Against Zinc-Induced Synaptic Impairment and Oxidative Stress Through the ERK1/2 Pathway. Frontiers in pharmacology 2022, 13:821521.

37. Tang Z, Lai CC, Luo J, Ding YT, Chen Q, Guan ZZ: Mangiferin prevents the impairment of mitochondrial dynamics and an increase in oxidative stress caused by excessive fluoride in SH-SY5Y cells. Journal of biochemical and molecular toxicology 2021, 35(4):e22705.

38. Tang Z, Guo M, Peng Y, Zhang T, Xiao Y, Ni R, Qi X: Quercetin reduces APP expression, oxidative stress and mitochondrial dysfunction in the N2a/APPswe cells via ERK1/2 and AKT pathways. bioRxiv 2022:2022.2009.2018.508406.

39. Li J, Xiang H, Huang C, Lu J: Pharmacological Actions of Myricetin in the Nervous System: A Comprehensive Review of Preclinical Studies in Animals and Cell Models. Front Pharmacol 2021, 12:797298.

40. Lai C, Chen Q, Ding Y, Su S, Liu H, Ni R, Tang Z: Rapamycin attenuated zinc-induced tau phosphorylation and oxidative stress in animal model: Involvement of dual mTOR/p70S6K and Nrf2/HO-1 pathways. bioRxiv 2021.

41. Puzzo D, Piacentini R, Fá M, Gulisano W, Li Puma DD, Staniszewski A, Zhang H, Tropea MR, Cocco S, Palmeri A et al: LTP and memory impairment caused by extracellular Aβ and Tau oligomers is APP-dependent. eLife 2017, 6.

42. Ono K, Li L, Takamura Y, Yoshiike Y, Zhu L, Han F, Mao X, Ikeda T, Takasaki J-i, Nishijo H et al: Phenolic Compounds Prevent Amyloid β-Protein Oligomerization and Synaptic Dysfunction by Site-specific Binding*. Journal of Biological Chemistry 2012, 287(18):14631–14643.

43. Alexiou A, Soursou G, Chatzichronis S, Gasparatos E, Kamal MA, Yarla NS, Perveen A, Barreto GE, Ashraf GM: Role of GTPases in the Regulation of Mitochondrial Dynamics in Alzheimer’s Disease and CNS-Related Disorders. Molecular neurobiology 2019, 56(6):4530–4538.

44. Manczak M, Kandimalla R, Yin X, Reddy PH: Hippocampal mutant APP and amyloid beta-induced cognitive decline, dendritic spine loss, defective autophagy, mitophagy and mitochondrial abnormalities in a mouse model of Alzheimer’s disease. Human molecular genetics 2018, 27(8):1332–1342.

45. Lee A, Kondapalli C, Virga DM, Lewis TL, Jr., Koo SY, Ashok A, Mairet-Coello G, Herzig S, Foretz M, Viollet B et al: Aβ42 oligomers trigger synaptic loss through CAMKK2-AMPK-dependent effectors coordinating mitochondrial fission and mitophagy. Nat Commun 2022, 13(1):4444.

46. Taguchi N, Ishihara N, Jofuku A, Oka T, Mihara K: Mitotic phosphorylation of dynamin-related GTPase Drp1 participates in mitochondrial fission. J Biol Chem 2007, 282(15):11521–11529.

47. Hung CH, Cheng SS, Cheung YT, Wuwongse S, Zhang NQ, Ho YS, Lee SM, Chang RC: A reciprocal relationship between reactive oxygen species and mitochondrial dynamics in neurodegeneration. Redox Biol 2018, 14:7–19.

48. Kim HY, Park SY, Choung SY: Enhancing effects of myricetin on the osteogenic differentiation of human periodontal ligament stem cells via BMP-2/Smad and ERK/JNK/p38 mitogen-activated protein kinase signaling pathway. European journal of pharmacology 2018, 834:84–91.

49. Zhao M, Wang Y, Li L, Liu S, Wang C, Yuan Y, Yang G, Chen Y, Cheng J, Lu Y et al: Mitochondrial ROS promote mitochondrial dysfunction and inflammation in ischemic acute kidney injury by disrupting TFAM-mediated mtDNA maintenance. Theranostics 2021, 11(4):1845–1863.

50. Jiang S, Nandy P, Wang W, Ma X, Hsia J, Wang C, Wang Z, Niu M, Siedlak SL, Torres S et al: Mfn2 ablation causes an oxidative stress response and eventual neuronal death in the hippocampus and cortex. Mol Neurodegener 2018, 13(1):5.

51. Jung Y, Shin SY, Lee YH, Lim Y: Flavones with inhibitory effects on glycogen synthase kinase 3β. Applied Biological Chemistry 2017, 60(3):227–232.

52. Nagaraj S, Want A, Laskowska-Kaszub K, Fesiuk A, Vaz S, Logarinho E, Wojda U: Candidate Alzheimer’s Disease Biomarker miR-483-5p Lowers TAU Phosphorylation by Direct ERK1/2 Repression. International journal of molecular sciences 2021, 22(7).

53. Mai M, Guo X, Huang Y, Zhang W, Xu Y, Zhang Y, Bai X, Wu J, Zu H: DHCR24 Knockdown Induces Tau Hyperphosphorylation at Thr181, Ser199, Ser262, and Ser396 Sites via Activation of the Lipid Raft-Dependent Ras/MEK/ERK Signaling Pathway in C8D1A Astrocytes. Molecular neurobiology 2022, 59(9):5856–5873.

54. Iroegbu JD, Ijomone OK, Femi-Akinlosotu OM, Ijomone OM: ERK/MAPK signalling in the developing brain: Perturbations and consequences. Neuroscience and biobehavioral reviews 2021, 131:792–805.

55. Foidl BM, Humpel C: Differential Hyperphosphorylation of Tau-S199, -T231 and - S396 in Organotypic Brain Slices of Alzheimer Mice. A Model to Study Early Tau Hyperphosphorylation Using Okadaic Acid. Frontiers in aging neuroscience 2018, 10:113.

56. Yang W, Liu Y, Xu QQ, Xian YF, Lin ZX: Sulforaphene Ameliorates Neuroinflammation and Hyperphosphorylated Tau Protein via Regulating the PI3K/Akt/GSK-3β Pathway in Experimental Models of Alzheimer’s Disease. Oxidative medicine and cellular longevity 2020, 2020:4754195.

57. Leroy K, Yilmaz Z, Brion JP: Increased level of active GSK-3beta in Alzheimer’s disease and accumulation in argyrophilic grains and in neurones at different stages of neurofibrillary degeneration. Neuropathology and applied neurobiology 2007, 33(1):43–55.

