## Supplemental Table 1 for "Myricetin protected against Aβ oligomer-induced synaptic impairment, mitochondrial function and oxidative stress in SH-SY5Y cells via ERK1/2/GSK-3β pathways"

**Supplemental Table 1. Antibodies used in this study.**

| **Antibody** | **Host** | **Specificity** | **WB/IF**  **Dilution** | | **Sources** | **Catalog No** |
| --- | --- | --- | --- | --- | --- | --- |
| Anti-beta Amyloid 1-42 | r | - | 1:1000 | | Abcam | #ab180956 |
| Anti-Tau S396 | r | p-Tau | 1:1000 | | Abcam | #ab32057 |
| Anti-Tau5 | m | - | 1:1000 | | Abcam | #ab80579 |
| SNAP25 | r | - | 1:5000 | | Abcam | #ab5666 |
| synaptophysin | r | - | 1:10000 | | Abcam | #ab32127 |
| PSD95 | r | - | 1:2000 | | Abcam | #ab18258 |
| Anti-p44/42 MAPK(Erk1/2)(137F5) | r | T-ERK1/2 | 1:1000 | | Cell Signaling | #4695 |
| Anti-P-p44/42 MAPK(Erk1/2)(Thr202/Tyr204)(D13.14.4E) | r | P-ERK1/2 | 1:2000 | | Cell Signaling | #4370 |
| Anti-GSK-3β | r | T-GSK-3β | 1:1000 | | Cell Signaling | #12456 |
| Anti-p-GSK-3β | r | P-GSK-3β | 1:1000 | | Cell Signaling | #9323 |
| Anti-Mitofusin 1 | r | Mfn1 | 1:1000 | | Abcam | #ab221661 |
| Anti-Mitofusin 2 | r | Mfn2 | 1:1000 | | Abcam | #ab124773 |
| Anti-Dynamin-related protein 1 | r | Drp1 | 1:1000 | | Abcam | #ab184247 |
| Anti-β-tubulin | m | - | 1:5000 | | Beyotime | #21463 |
| Anti-β-actin | m | - | 1:25000 | | Affinity Biosciences | #T0022 |
| Anti-4-Hydroxynonenal | r | 4-HNE | 1:200 | Abcam | | #ab46545 |
| Anti-8-Hydroxy-2'-deoxyguanosine | m | 8-OHdG | 1:200 | Abcam | | #ab48508 |
| Anti-rabbit IgG (H+L) (HRP) | g | - | 1:5000 | Thermo Fisher | | #32260 |
| Anti-mouse IgG (H+L) (HRP) | g | - | 1:5000 | Thermo Fisher | | #32230 |
| Anti-Mouse IgG (H+L) Cyanine3 | g | - | 1:200 | Thermo Fisher | | #31188 |
| Anti-Rabbit IgG (H+L) Alexa Fluor™546 | d | - | 1:200 | Thermo Fisher | | #31466 |

T: total; P: phosphorylated; r: rabbit; m: mouse; g: goat; d: donkey; IF: immunofluorescence; WB: western blot;
