## Supplemental Table 2 for "Myricetin protected against Aβ oligomer-induced synaptic impairment, mitochondrial function and oxidative stress in SH-SY5Y cells via ERK1/2/GSK-3β pathways"

**Supplemental Table 2. Chemicals and kits used in this study.**

| **Chemical/kit** | **Sources** | **Catalog No** |
| --- | --- | --- |
| Myricetin | MedChemExpress | #HY-15097 |
| β-Amyloid (1-42) | GL Biochem (shanghai) Co., Ltd | #GLS-52487 |
| SH-SY5Y cell | ATCC | # CRL-2266 |
| Cell Counting Kit-8 | MedChemExpress | # HY-K0301 |
| DAPI solution | Solarbio | # C0065 |
| RIPA buffer(high) | Solarbio | #R0010 |
| Protease inhibitor mixture | Solarbio | #P6730 |
| Fetal Bovine Serum | Thermo Fisher | #10099141C |
| HFIP | Sigma-Aldrich | #105228 |
| DMSO | Sigma-Aldrich | #D2650 |
| PEG 300 | Sigma-Aldrich | #91462 |
| Tween-80 | Sigma-Aldrich | #P1754 |
| Triton™ X-100 | Sigma-Aldrich | #X100 |
| Pierce™ BCA Protein Test Kit | Thermo Fisher | #23227 |
| ROS Assay Kit | Beyotime | #S0033S |
| Mitochondrial membrane potential assay kit with JC-1 | Beyotime | #C2006 |
